# Combinatorial viral vector-based and live-attenuated vaccines without an adjuvant to generate broader immune responses to effectively combat pneumonic plague

**DOI:** 10.1101/2021.10.28.466384

**Authors:** Paul B. Kilgore, Jian Sha, Emily K. Hendrix, Vladimir L. Motin, Ashok K. Chopra

## Abstract

Mice immunized with a combination of an adenovirus vector (Ad5-YFV) and live-attenuated (LMA)-based vaccines were evaluated for protective efficacy against pneumonic plague. While the Ad5-YFV vaccine harbors a fusion cassette of three genes encoding <u>Y</u>scF, <u>F</u>1, and <u>L</u>crV, LMA represents a mutant of parental *Yersinia pestis* CO92 deleted for genes encoding <u>L</u>pp, <u>M</u>sbB, and <u>A</u>il. Ad5-YFV and LMA were either administered simultaneously (1-dose regimen) or 21 days apart in various order and route of administration combinations (2-dose regimen). The 2-dose regimen induced robust immune responses to provide full protection to animals against parental CO92 and its isogenic F1 (CAF^-^)-deletion mutant challenges during both short- and long-term studies. Mice intranasally (i.n.) immunized with Ad5-YFV first followed by LMA (i.n. or intramuscularly [i.m.]) had higher T- and B-cell proliferative responses and LcrV antibody titers than those in mice vaccinated with LMA (i.n. or i.m.) first ahead of Ad5-YFV (i.n.) during the long-term study. Specifically, the needle- and adjuvant-free vaccine combination (i.n.) is ideal for use in plague endemic regions. Conversely, with a 1-dose regimen, mice vaccinated with Ad5-YFV i.n. and LMA by the i.m. route provided complete protection to animals against CO92 and its CAF^-^ mutant challenges, and elicited Th1/Th2, as well as Th17 responses, thus suitable for emergency vaccination during a plague outbreak or bioterrorist attack. This is a first study in which a viral vector-based and live-attenuated vaccines were effectively used in combination representing adjuvant- and/or needle- free immunization, with each vaccine triggering a distinct cellular immune response.

**Importance:** *Yersinia pestis*, the causative agent of plague, is a Tier-1 select agent and a re-emerging human pathogen. A 2017 outbreak in Madagascar with >75% of cases being pneumonic and 8.6% causalities emphasized the importance of the disease. The World Health Organization has indicated an urgent need to develop new generation subunit and live-attenuated plague vaccines. We have developed a subunit vaccine including three components (<u>Y</u>scF, <u>F</u>1, and Lcr<u>V</u>) using an adenovirus platform (Ad5-YFV). In addition, we have deleted virulence genes of *Y. pestis* (*e.g., lpp*, *<u>m</u>sbB*, and *<u>a</u>il*) to develop a live-attenuated vaccine (LMA). Both of these vaccines generated robust humoral and cellular immunity and were highly efficacious in several animal models. We hypothesized the use of a heterologous prime-boost strategy or administrating both vaccines simultaneously could provide an adjuvant- and/or a needle- free vaccine(s) which have attributes of both vaccines for use in endemic regions and during an emergency situation.

## Introduction

The zoonotic bacterium *Yersinia pestis*, the causative agent of bubonic and pneumonic plague, is a Tier-1 select agent. Historically, three plague pandemics resulted in more than 200 million deaths worldwide (1). The endemic nature of the disease has resulted in an ever-increasing number of human plague cases worldwide, including the U.S., with a case fatality rate of 18% (2). In the most recent major outbreak of plague in Madagascar (2017), although millions of doses of levofloxacin were prescribed to treat people suspected of contracting plague, the mortality rate still was at 8.6% (3). Notably, in this outbreak, >75% of the cases were pneumonic in nature in contrast to traditional bubonic plague, the latter of which has a slower disease course and a longer window for effective antibiotic treatment (4, 5). However, for pneumonic plague, antibiotic therapy must be initiated within 24 hours of the onset of symptoms to be effective, and if left untreated, fatality rate is almost 100% (6). Therefore, vaccination is the most effective way to prevent plague. Unfortunately, there are no Food and Drug Administration (FDA)-approved plague vaccines (7), and those currently under clinical trials are all based on two antigens: F1 (capsular antigen) and LcrV (low calcium response V antigen, a type 3 secretion system [T3SS] component) (8–10). Consequently, in individuals vaccinated with the F1-V-based vaccines and infected with the F1-negative strain of *Y. pestis*, protection would be reliant on only immune responses generated against LcrV. Indeed, a LcrV-based vaccine alone did not provide complete protection against pneumonic plague against F1-negative *Y. pestis* challenge, and the F1-V-based vaccines were much less effective against bubonic and pneumonic plague upon challenge with the F1-negative *Y. pestis* strain C12 (11–13). Similarly, the F1-V-based vaccines were not completely protective against pneumonic plague in African green monkeys (14). More alarming is the existence of LcrV polymorphism in *Y. pestis* as well as variants in *Y. enterocolitca* and *Y. pseudotuberculosis* strains with little cross protection (15–18). Therefore, it is necessary to develop new plague vaccines and evaluate different immunization strategies that could be effective against the F1-negative *Y. pestis* strains, those which harbor LcrV variants, as well as against human population with varying immune responses in endemic regions or during an outbreak/biothreat situation.

In our laboratory, we have developed two types of plague vaccine candidates, namely LMA and Ad5-YFV. The live-attenuated vaccine LMA is a triple deletion mutant of *Y. pestis* CO92 in which genes encoding Braun lipoprotein (Lpp), an acyltransferase (MsbB), and the attachment invasion locus (Ail), were deleted (19). While Lpp activates pro-inflammatory cascade by binding to toll-like receptor 2 (TLR-2) (20, 21), MsbB adds lauric acid to the lipid A moiety of lipopolysaccharide (LPS), which triggers TLR-4 signaling (22, 23). Ail, in addition to promoting attachment and invasion of *Y. pestis* to the host, provides serum resistance to the organism (19, 24).

The Ad5-YFV is a replication deficient adenovirus type 5 vector-based vaccine containing genes for three plague antigens: fraction 1 capsule-like antigen F1, tip protein of the T3SS LcrV, and YscF that forms the barrel structure of T3SS needle (25). Using a prime-boost immunization regimen (21 days apart), both vaccines when used individually, elicited robust humoral and cell-mediated immune responses in animals and conferred 100% protection against lethal *Y. pestis* CO92 challenge (26, 27). Importantly, each of these vaccines has its own unique characteristics. Notably, we observed a significant induction of CD4^+^ IL-17^+^-producing T-cells in LMA vaccine-immunized mice (27, 28). IL-17 production is an important correlate of protection against plague in the absence of protective antibodies (29, 30). However, such a T-cell population was not detected in animals immunized with the Ad5-YFV vaccine (26). Further, the Th1 immune response was favored after vaccination of mice with the Ad5-YFV vaccine over Th2, possibly due to the Ad5 vector used, while the Th2 immune response was favored after immunization of animals with the LMA vaccine (26, 28). In our past studies, the Ad5-YFV vaccine was always delivered intranasally (i.n.), while the LMA vaccine was administered by the i.m. or the i.n. route (19, 25–28).

Like other live-attenuated vaccines, one of the major advantages of LMA is that it delivers a large array of antigens in their native state, closely mimicking natural infection, and such vaccines are expected to provide protection against all circulating *Y. pestis* variants in the nature (30–34). While the viral vector-based vaccines, especially the replication deficient ones such as Ad5-YFV, are much safer than the live-attenuated vaccines in general, both types of vaccines have their own disadvantages. The most obvious limitation of live-attenuated vaccines is the safety profile, especially in immunocompromised individuals, and the potential concern for reversion (35–37). However, the LMA mutant was rationally designed with complete deletion of three genes located at different locations on the bacterial genome (19). Because of its high level of virulence attenuation and rapid clearance (within 12-24 h) while retaining immunogenicity in animals (27), LMA vaccine was excluded from the Centers for Disease Control and Prevention (CDC) select agent list (https://www.selectagents.gov/sat/exclusions/hhs.html).

The Ad5-YFV vaccine only incorporates three plague antigens: F1, LcrV and YscF, and the F1-minus strains of *Y. pestis* have been isolated from humans/animals and are as virulent as the wild-type (WT) plague bacterium (38, 39). In addition, hypervariable regions within the LcrV protein have been described and antibody responses to these LcrV variants are not cross-protective (18). Therefore, we have successfully added a third protective antigen YscF in this vaccine to circumvent disadvantages of F1-V-based vaccines (25, 40). However, it is still plausible that the Ad5-YFV vaccine alone may not be efficacious against all circulating *Y. pestis* variants, although it is shown to be protective (100%) against CAF^-^ mutant of CO92 (26). Importantly, both of our vaccines do not require an adjuvant to boost immune responses, unlike F1-V-based vaccines which employ Alum (8, 9, 11, 41–43).

To alleviate some of the above concerns, in this study, we carried out a heterologous immunization strategy in which both LMA and Ad5-YFV vaccines were administered either simultaneously (1-dose regimen) or in a prime-boost format (2-dose regimen). Our results showed almost all the heterologous vaccination groups of mice induced robust immune responses and provided full protection against lethal challenge doses of both parental and CAF^-^ mutant of *Y. pestis* CO92; albeit using potentially slightly different mechanisms. This is the first detailed plague vaccine study with heterologous vaccination regimens that involved a live-attenuated vaccine LMA and a viral vector-based vaccine Ad5-YFV.

## Results

### Virulence and immunogenic characterization of the LMA vaccine candidate in iron over loaded conventional and or in immunocompromised mice

We have previously shown LMA mutant to be highly attenuated in conventional (immunocompetent) mice (19, 27). To further evaluate its attenuation, we tested safety of the LMA mutant during iron overload conditions in conventional mice to mimic hemochromatosis. We also tested its safety in immunocompromised Rag1 KO mice.

We demonstrated that all of the mice challenged with the LMA vaccine (5×10^6^ CFU, more than double the vaccination dose) by the i.n. route survived irrespective of whether the animals were iron-overloaded or not (**Fig. 1A**) with a minimal loss in the body weight (**Fig. 1B**). On the contrary, mice infected with the same dose of the KIM/D27 strain died by day 5 in the presence of FeCl_2_, while 80% of the animals succumbed to infection without iron-overload (**Fig. 1A**). At a lower challenge dose of 2×10^5^ CFU, 80% of the iron-overloaded mice succumbed by day 7 while only 20% of the non-iron-overloaded mice succumbed (**Fig. 1A**). In contrast to the LMA mutant, KIM/D27 strain-infected mice showed a dramatic loss in the body weight even at the lower challenge dose of 2×10^5^ CFU (**Fig. 1B**).

**Figure 1:**
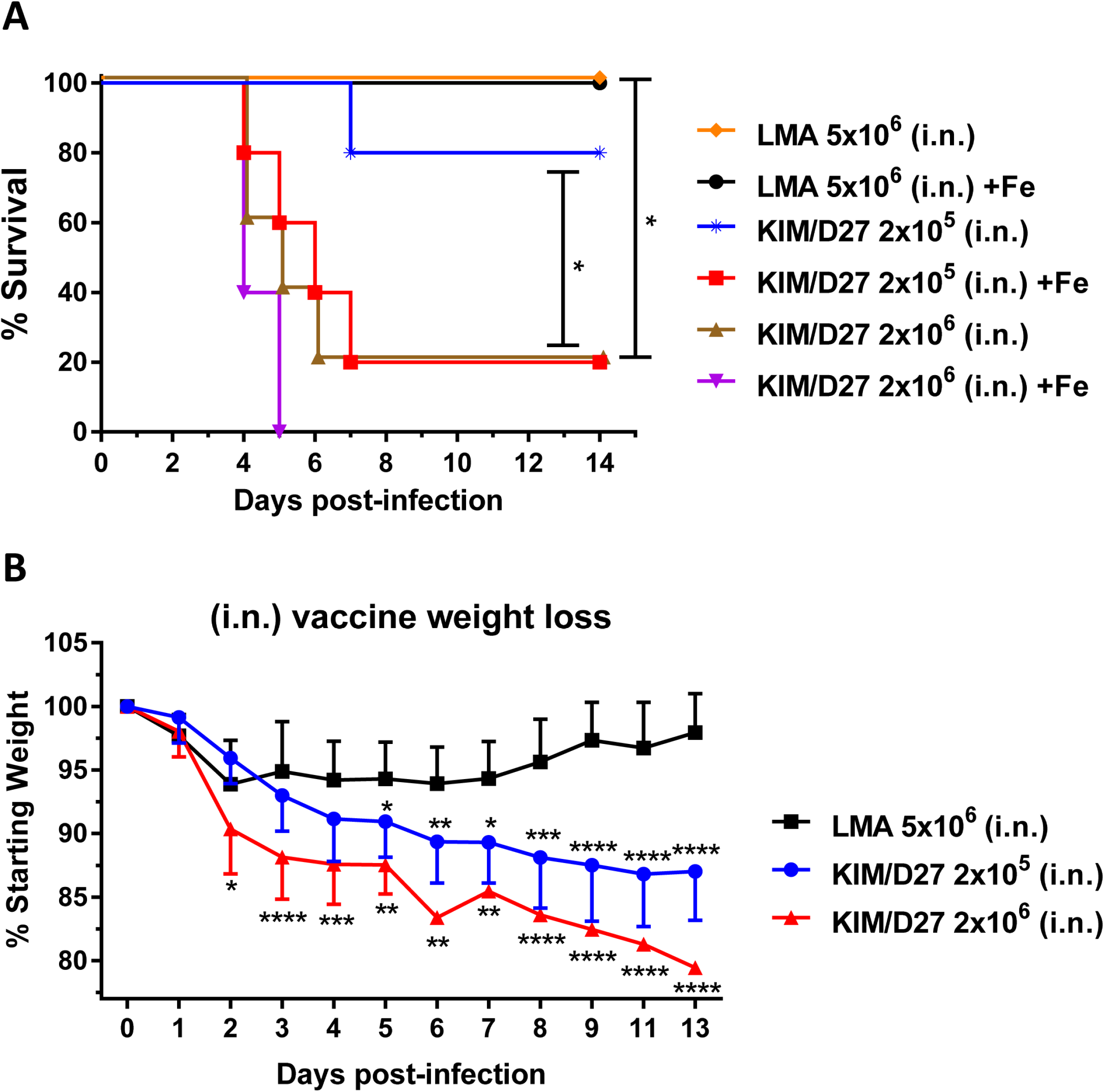
Iron-overload condition does not restore virulence of the LMA vaccine. Swiss-Webster mice (n=5/group) were injected with 75µg of FeCL_2_·4H_2_O 1h prior to infection and then infected with either the LMA vaccine or the KIM/D27 strain. Loss in body weight and mortality were monitored for 14 days; animals without iron-overload were used as controls (**A and B**). Kaplan-Meier analysis with log-rank (Mantel-Cox) test was used for analysis of animal survival. Two-way ANOVA with Tukey’s post hoc test was used to calculate significant differences in body weight loss between vaccine groups. Asterisks represent the statistical significance between two groups indicated by a line for (**A**) and between LMA and the indicated dose of KIM/D27 in (**B**). Since absence or presence of iron did not affect animal body weight, combined data were plotted. Two biological replicates were performed. * p<0.05, **p<0.01, *** p<0.001 **** p<0.0001.

We then infected Rag1 KO mice with 4 LD_50_ of CO92 by either the i.n. or the i.m. route since these two routes were previously used for vaccination studies with LMA (19, 27, 28). As shown in **Fig. S1A**, 100% of mice infected by the i.n. or the i.m. route succumbed to infection by day 4-5 p.i., with up to ∼20% body weight loss by day 4. We then examined bacterial burden in the lungs, liver, and spleen of these mice and found greater than 10^8^ CFU of *Y. pestis* in both the lungs and the spleen, while a significantly higher *Y. pestis* burden, with up to a more than 10^9^ CFU, was observed in the liver (**Fig. S1B**).

We then evaluated the LMA mutant, when delivered at a vaccination dose of 2.0 x 10^6^ CFU by both i.n. and i.m. routes, in Rag1 KO mice. Five days p.i., five mice infected with LMA from each infection route were necropsied and examined for the presence of the mutant either at the infection site (lungs or muscle) or in the spleen as the result of systematic dissemination. As shown in **Fig. 2A**, there was no detectable LMA mutant in the spleen of mice infected by either the i.n. or the i.m. route. We also did not detect any LMA mutant at the injection site of muscle. We did enumerate 300 CFU in the lungs of one mouse that was infected by the i.n. route; however, this number was much lower than the initial infection dose of 2.0 x 10^6^ CFU. Further, no LMA mutant was detected in the lungs of other mice infected by the i.n. route (**Fig. 2A**). This was also indicated by the fact that all the mice infected with 2.0 x 10^6^ CFU of LMA survived up to 28 days p.i. (**Fig. 2B**) without any clinical signs of the disease and a minimal body weight loss similar to that shown in conventional mice (**Fig. 1B)**.

**Figure 2:**
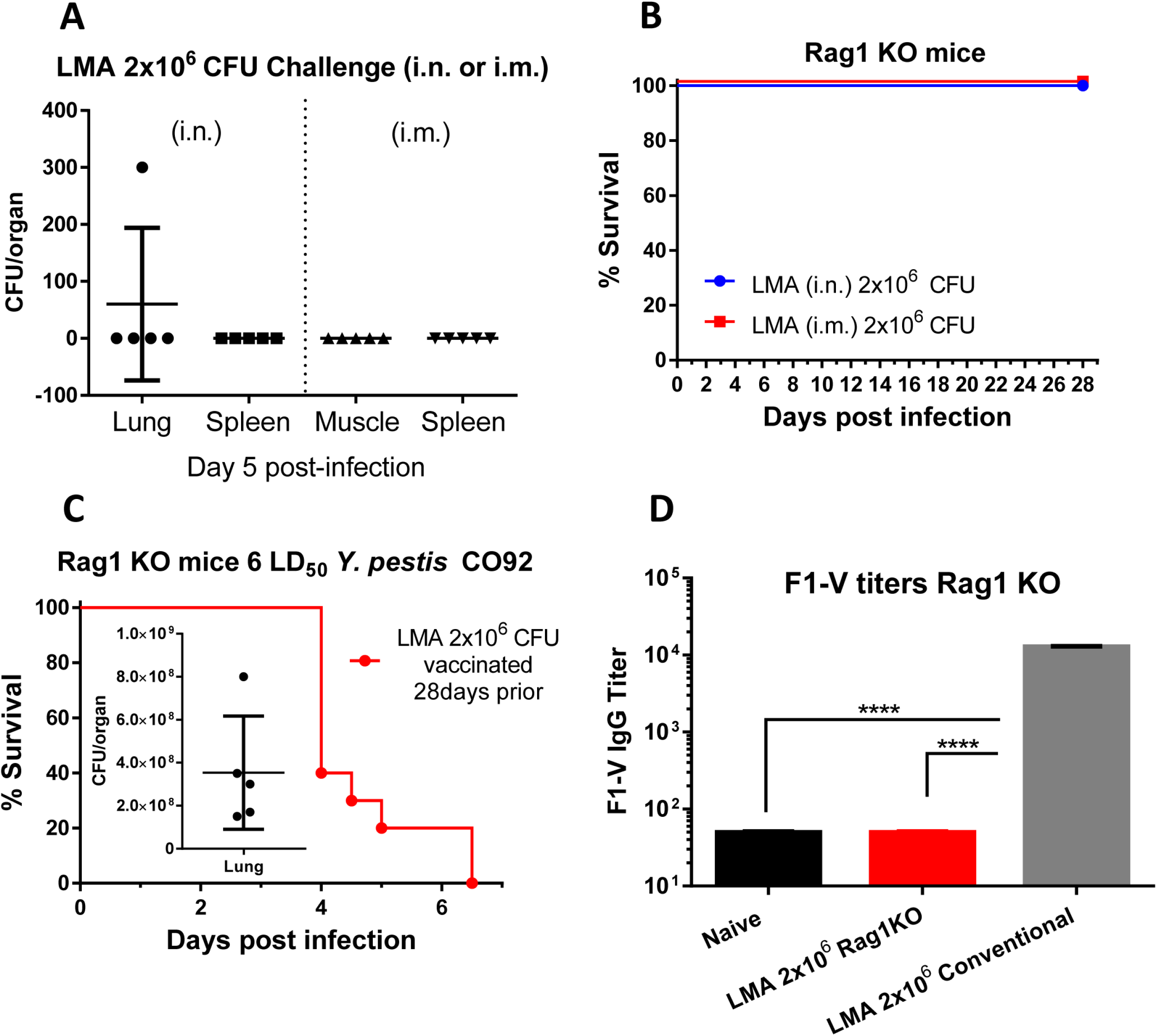
Attenuation and immunologic characterization of LMA vaccine in Rag1 KO mice. C57BL6 Rag1 KO mice were infected with 2.0 x 10^6^ CFU of LMA by either the i.n. or the i.m. route (n=10 each route). On day 5 p.i., 5 animals from each infection route were euthanized and spleen, lungs or muscle (depending on the infection route) were collected to quantify the number of LMA (**A**). The survival of the remaining LMA infected animals (5 from each route) were monitored for up to 28 days p.i. (**B**). After 28 days of LMA infection, all surviving mice (n=10) were challenged with 6 LD_50_ of CO92. The mortality of animals was recorded and bacterial loads in the lungs from 5 moribund animals (inset) were enumerated (**C**). F1-V specific IgG titers were evaluated by ELISA from sera collected at day 3 post CO92 challenge. Sera collected from uninfected naïve Rag1 KO mice and conventional mice immunized with LMA (2.0 x 10^6^ CFU i.m.) served as negative and positive controls, respectively (**D**). One-way ANOVA was used to determine significance between groups for bacterial burdens and antibody titers. Kaplan-Meier analysis with log-rank (Mantel-Cox) test was used for analysis of animal survival. Asterisks represent the statistical significance between two groups indicated by a line and *** p<0.001. Two biological replicates were performed, and data plotted.

These surviving mice were then challenged with 6 LD_50_ of CO92. As expected, all LMA infected Rag1 KO mice succumbed to CO92 challenge with an overall >10^8^ CFU of *Y. pestis* CO92 present in the lungs (**Fig. 2C**) with ∼15% loss in body weight over time. Further, there were no F1-V specific IgG antibodies in the sera of LMA (pooled from i.n. and i.m. infected) Rag1 KO mice on day 3 post CO92 challenge as compared to naïve Rag1 KO mice. However, a significantly higher level of F1-V IgG antibodies was generated in the LMA-immunized conventional (immunocompetent) mice (pooled sera from i.n. and i.m. infected) from a parallel independent study (**Fig. 2D**).

### Strong immune responses were elicited in conventional mice by heterologous vaccination with either a 1- or 2-dose (prime-boost) regimen

Mice were immunized with Ad5-YFV and LMA vaccines in either a 1- or 2-dose regimen. In a 1-dose regimen, both Ad5-YFV and LMA vaccines were administered simultaneously, while in a 2-dose regimen, Ad5-YFV and LMA vaccines were delivered 21 days apart in various order and route combinations (**Fig. 3A**). The immunization schedule for either 1- or 2-dose regimens is depicted in **Fig. 3B**). Three weeks after completion of the vaccinations, the immunized and control mice were challenged with 100 LD_50_ of either CO92 or its F1 deletion mutant CAF^-^. As shown in **Fig. 3C** and **D**, all of 2-dose heterologous prime-boost vaccinated mice, regardless of the order in which the vaccines were administered or the route of vaccination by which LMA was delivered, were 100% protected from both CO92 and its CAF^-^ strain challenge with no body weight loss and other clinical symptoms of the disease. A 100% protection was also observed for mice simultaneously immunized with Ad5-YFV (i.n.) and LMA (i.m) during CO92 and CAF^-^ challenges. However, when both Ad5-YFV and LMA vaccines were simultaneously administered i.n., the immunized mice had 50% (during CAF^-^ strain challenge) and 63% (during CO92 challenge) survival rates (**Fig. 3E and F**).

**Figure 3:**
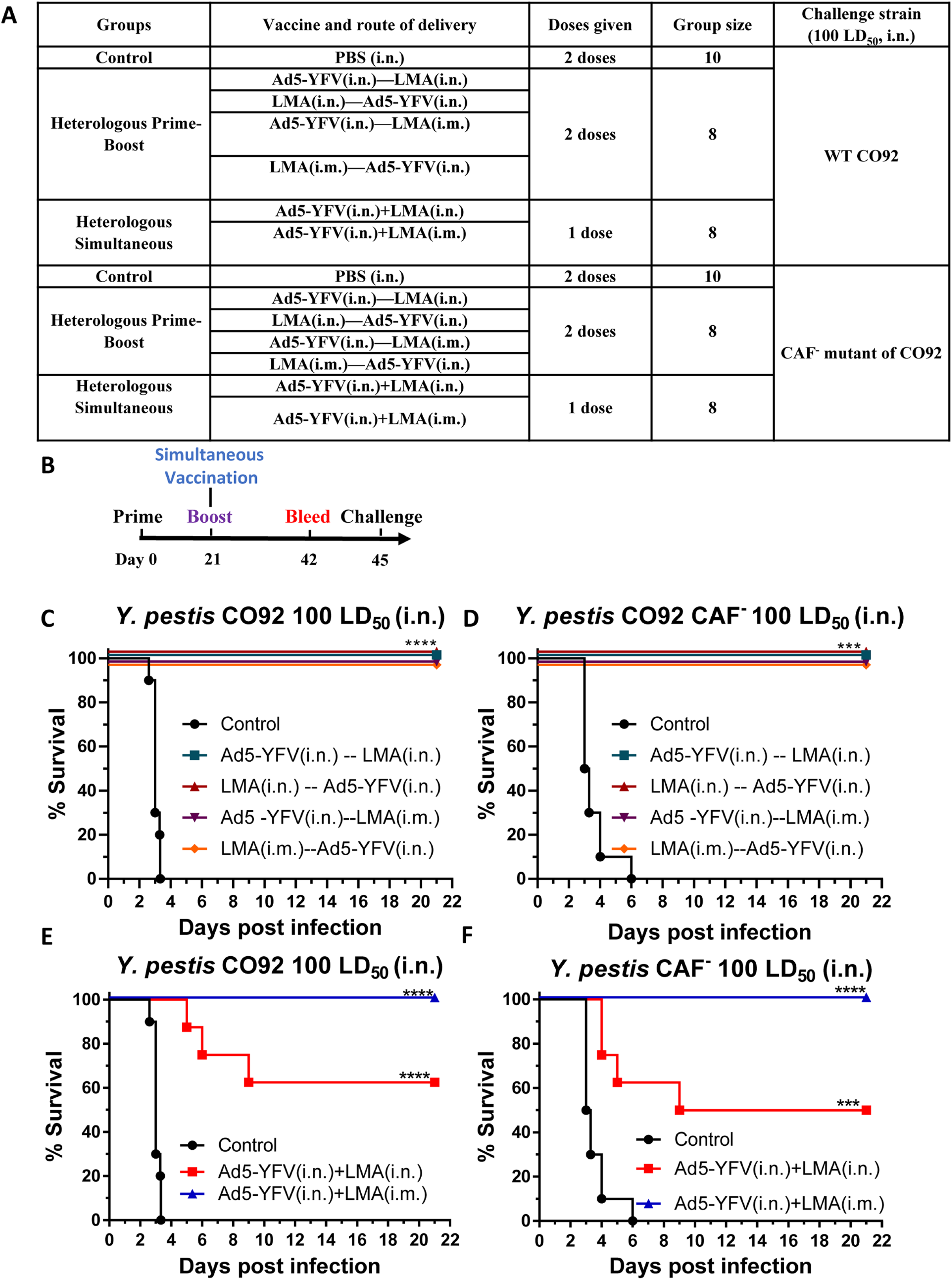
Short-term heterologous vaccination study with conventional mice. Mice (n=8-10 per group) were immunized heterologously with Ad5-YFV and LMA vaccines in either a 1- or a 2-dose regimen. In 1-dose regimen, both Ad5-YFV and LMA vaccines were delivered simultaneously, while in a 2-dose regimen, Ad5-YFV and LMA vaccines were administered 21 days apart in various order and combinations. Mice receiving PBS were used as controls. The composition of the groups and the experiment time course are depicted in panels **A** and **B** respectively. Three weeks after completion of the vaccination schedule, mice were challenged with 100 LD_50_ of either CO92 (**C** and **E**) or its F1 deletion mutant CAF^-^ (**D** and **F**) and observed for morbidity and mortality for 21 days. Kaplan-Meier analysis with log-rank (Mantel-Cox) test was used for analysis of animal survivals. Asterisks represent the statistical significance between the indicated groups to the naïve control mice. *** p<0.001, ****p<0.0001. Two biological replicates were performed, and data plotted.

We then measured IgG antibody titers to rF1-V fusion protein in sera of immunized mice collected on day 42 prior to the challenges. In general, all vaccinated groups of animals had notable increases in antibody titers which were 2-3 logs higher compared to that of the naive controls (**Fig. 4A**). Among the immunized mice, relatively lower F1-V antibody titers were noted in animals immunized with either a 2-dose regimen of Ad5-YFV (i.n.)-LMA (i.m) or a 1-dose regimen of Ad5-YFV (i.n.) + LMA (i.n.). However, a significant difference was only observed between the 2 simultaneously immunized groups of mice (**Fig. 4A**, red versus blue bars).

**Figure 4:**
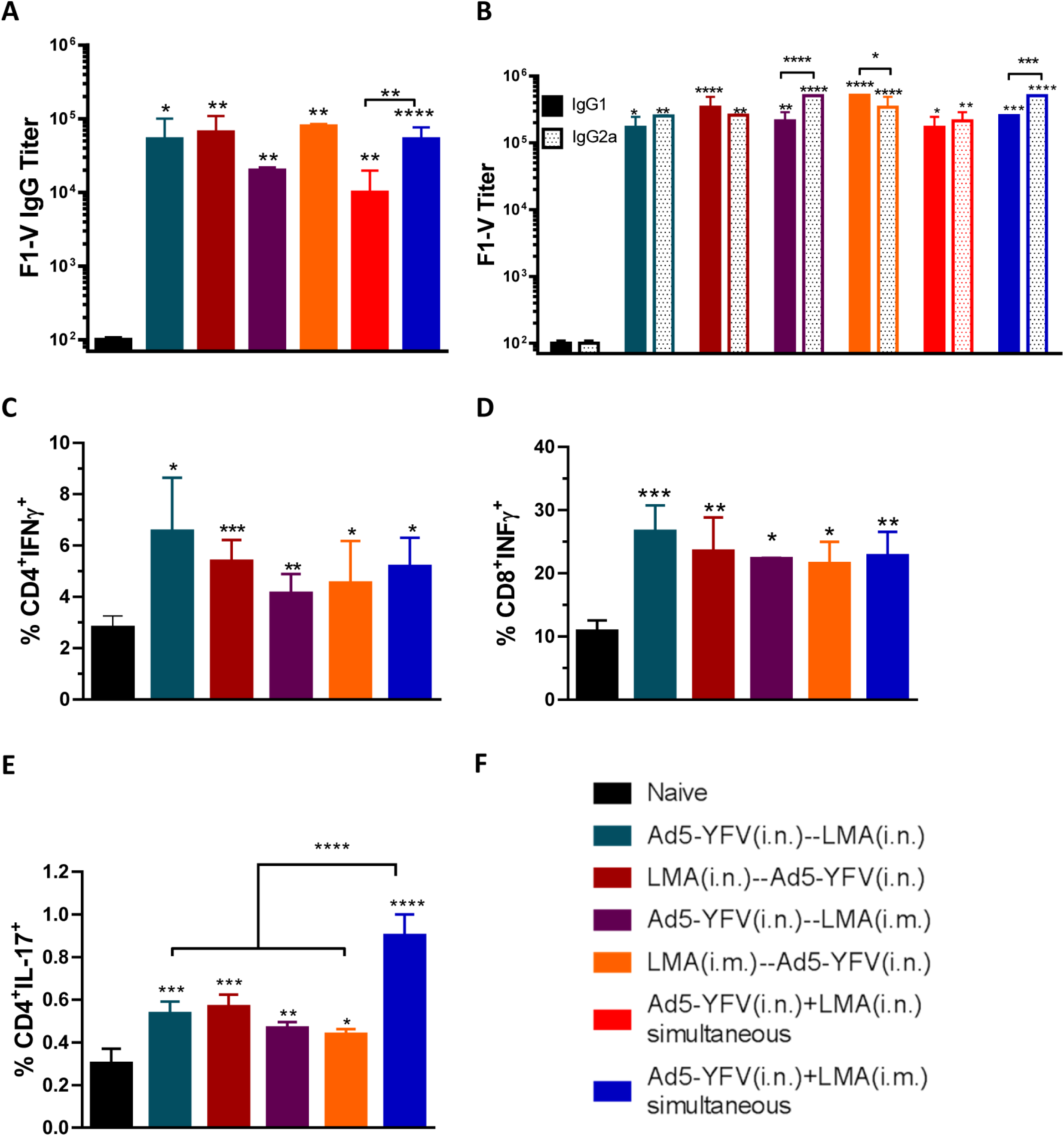
Humoral and cell-mediated immune responses during short-term heterologous vaccination study. Sera were collected 21 days after the last immunization from both immunized and naïve control mice as described in Fig. 3. ELISA was performed to evaluate specific F1-V total IgG titers (**A**) as well as its isotype IgG1 and IgG2a titers (**B**). In a separate experiment, mice (n=5) were similarly immunized as described in Fig. 3, however, the group in which mice were simultaneously immunized with LMA and Ad5-YFV i.n. was excluded. Twenty-one days after completion of the vaccination course, spleens were harvested. Splenocytes were isolated and stimulated with PMA, Ionomycin, and Brefeldin A. Subsequently, splenocytes were surface stained for CD3, CD4, and CD8 followed by intracellular staining for IFNγ (**C**,**D**) and IL-17A (**E**). Different groups of mice used are depicted in **F**. Statistical analysis was performed using One-way ANOVA with Tukey’s post-hoc test (**A**, **C**, **D**, **E**) or Two-way ANOVA with Tukey’s post-hoc test (**B**) to determine significance. Asterisks directly above bars indicated significance compared to control group while asterisks with comparison bars denoted significance between the indicated groups. *p<0.05, **p<0.01, ***p<0.001, ****p<0.0001. Two biological replicates were performed, and data plotted. *In vitro* studies had 3 replicates.

We also examined the isotypes of F1-V IgG antibodies to gauge Th1 versus Th2 bias. In general, all vaccinated groups of mice had significantly higher levels of F1-V specific IgG1 and IgG2a as compared to those of the naive animals. For simultaneously vaccinated groups of mice, there were generally higher levels of F1-V specific IgG2a over IgG1; however, a significant difference was only observed for animals that received Ad5-YFV (i.n.) and LMA (i.m.) (**Fig 4B**). For the 2-dose vaccinated groups of mice, when Ad5-YFV was administered first (teal and purple bars), there were always higher levels of IgG2a over IgG1, while the difference was only significant in the Ad5-YFV (i.n.)-LMA (i.m.) immunized group of mice as shown in purple bars (**Fig. 4B**). In contrast, when the LMA vaccine was delivered as the first dose (crimson and orange bars), higher levels of IgG1 over IgG2a were observed but again these differences were only significant when LMA was administered by the i.m. route as shown in orange bars (**Fig. 4B**).

We next examined cell-mediated immune responses in immunized mice from the groups which showed 100% protection during the *Y. pestis* challenges (**Fig. 3C-F**). In a separate experiment, splenocytes were isolated from vaccinated mice 21 days after the last vaccination dose and stimulated with PMA and Ionomycin. All vaccination groups had significantly higher populations of CD4^+^ IFNγ^+^, CD8^+^ IFNγ^+^ and CD4^+^ Il-17^+^cells than those of naive mice (**Fig 4C-E**). Among the vaccinated groups, mice i.n. immunized with either LMA or Ad5-YFV first in a 2-dose regimen (teal and crimson bars) showed slightly higher percentages of IFNγ producing CD4^+^ and CD8^+^ T cells than all other groups of immunized mice; however, no significant differences were observed (**Fig. 4C and D**).

On the other hand, notably higher population of IL-17 producing CD4^+^ T cell was noticed in the group of mice simultaneously immunized with Ad5-YFV (i.n.) and LMA (i.m.) as compared to all the 2-dose regimen immunized groups of mice (**Fig. 4E**, blue bar). Interestingly, although it did not reach significant level, mice i.n. immunized with either LMA or Ad5-YFV first in a 2-dose regimen (teal and crimson bars) showed slightly higher percentages of IL-17 producing CD4^+^ T cells than all other 2-dose regimen immunized groups of animals. A similar trend (to IL-17) was also observed for the IFNγ producing CD4^+^ and CD8^+^ T cells (**Fig. 4C and D**).

### Robust immune response was sustained in conventional mice with a 2-dose regimen vaccination during a long-term study

After examining the above survival data and immune responses from both 1- and 2-dose heterologous vaccination regimens, we chose to focus on the 2-dose regimen for further evaluation in a long-term vaccination study. The sole reason for this was to develop a needle-free vaccination protocol although comparisons were also made with the i.m. vaccinated animals. During the long-term study, we examined humoral and cell-mediated responses at 42 days after the 2^nd^ dose of vaccination as well as at 3 days post CO92 challenge. The details of immunization regimens and schedules of the study are shown in **Figs. 5A and B**.

**Figure 5:**
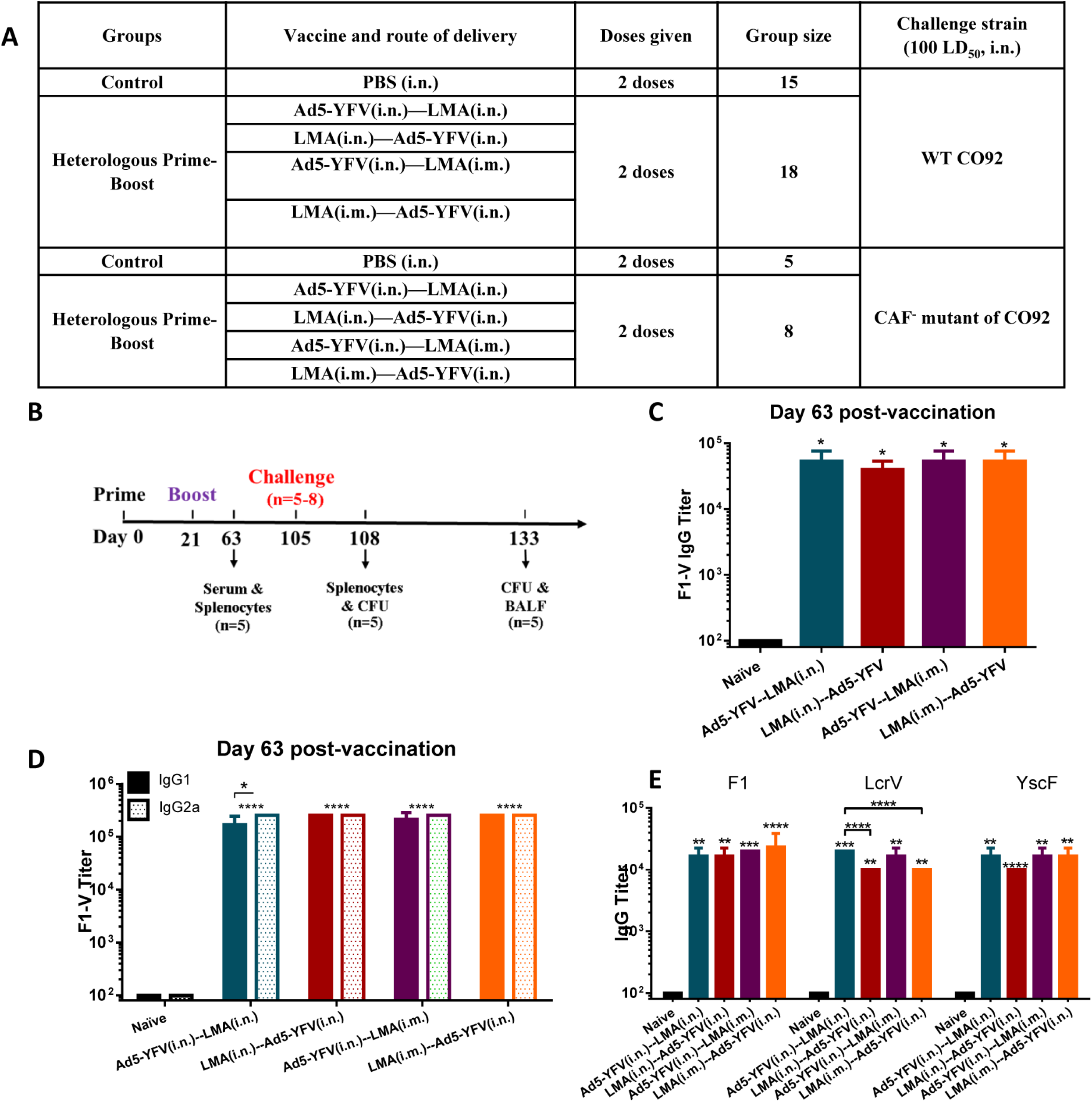
Antibody responses during long-term heterologous prime-boost vaccination study. Mice were immunized with Ad5-YFV and LMA vaccines in 2-dose (prime-boost) regimens in which Ad5-YFV and LMA were administered 21 days apart in various combinations. The composition of the groups and the experiment time course are depicted in panels **A** and **B,** respectively. Sera were collected on day 63 of the study which was 42 days after the 2^nd^ vaccination dose. The total IgG and its isotypes IgG1/IgG2a titers specific to F1-V were determined by ELISA and displayed in (**C**) and (**D**), respectively, while, the total IgG titers specific to individual antigens F1, LcrV, and YscF were shown in (**E**). Statistical significance was determined by One-way ANOVA with Tukey’s post-hoc test (**A**, **C**, **D**) and by Two-way ANOVA with Tukey’s post-hoc test (**B**). Asterisks directly above bars indicated significance compared to control group while asterisks with comparison bars denoted significance between the indicated groups. *p<0.05, **p<0.01, ***p<0.001, ****p<0.0001. Two biological replicates were performed, and data plotted. *In vitro* studies had 3 replicates.

As shown in **Fig. 5C**, a similar levels of F1-V specific antibodies were detected in the sera across all vaccinated groups of mice and were all significantly higher (>2 logs) than that of naïve control animals (**Fig. 5C**). Further, antibody isotype analysis revealed a generally balanced level of IgG1 and IgG2a in the immunized mice except for animals immunized first with Ad5-YFV (teal and purple bars) which had higher levels of IgG2a over IgG1. However, the difference was only significant in mice immunized with Ad5-YFV (i.n.)-LMA (i.n.) (**Fig. 5D**, teal bars).

To further dissect the antibodies elicited by the vaccination, we evaluated antibody titers to each individual plague antigen F1, LcrV, and YscF that were the components of Ad5-YFV vaccine. As shown in **Fig. 5E**, all immunized mice produced a similar level of IgG against each individual antigen and they were all significantly higher than that of naive control mice. To be more specific, antibody titers against F1 and YscF were comparable among all vaccinated groups of mice. However, antibody titers against LcrV were significantly higher in mice immunized with Ad5-YFV (i.n.)-LMA (i.n.) (teal bar) as compared to mice vaccinated with either LMA (i.n.)-Ad5-YFV (i.n.) (crimson bar) or LMA (i.m.)-Ad5-YFV (i.n.) (orange bar) (**Fig. 5E**).

The isolated splenocytes from mice 42 days post 2^nd^ dose of vaccine were stimulated with *Y. pestis* specific rF1-V fusion antigen (100 µg/ml) to induce cell proliferation by measuring incorporation of BrdU in the newly synthesized chromosomal DNA (44). As shown in **Fig. 6**, a significant T- and B-cell proliferation was generally noticed in all of the vaccinated groups as compared to the naïve controls. Interestingly, a much stronger T- and B-cell proliferation was achieved in mice which were first vaccinated with Ad5-YFV (teal and purple bars) than those mice which were first immunized with LMA (crimson and orange bars) regardless of the route by which LMA was administered (**Fig. 6**). It is plausible that administration of LMA first might have somewhat of a toxic effect on T- and B-cells, an effect not observed when Ad5-YFV was delivered first and could account for less T- and B-cell proliferation. Important to note is that antibody titers to LcrV (**Fig. 5E**) were also lower when LMA was used first followed by Ad5-YFV for vaccination of mice. We suspect that antibodies to YscF, the third component in the Ad5-YFV vaccine, compensates for this lower LcrV antibody titers in protection.

**Figure 6:**
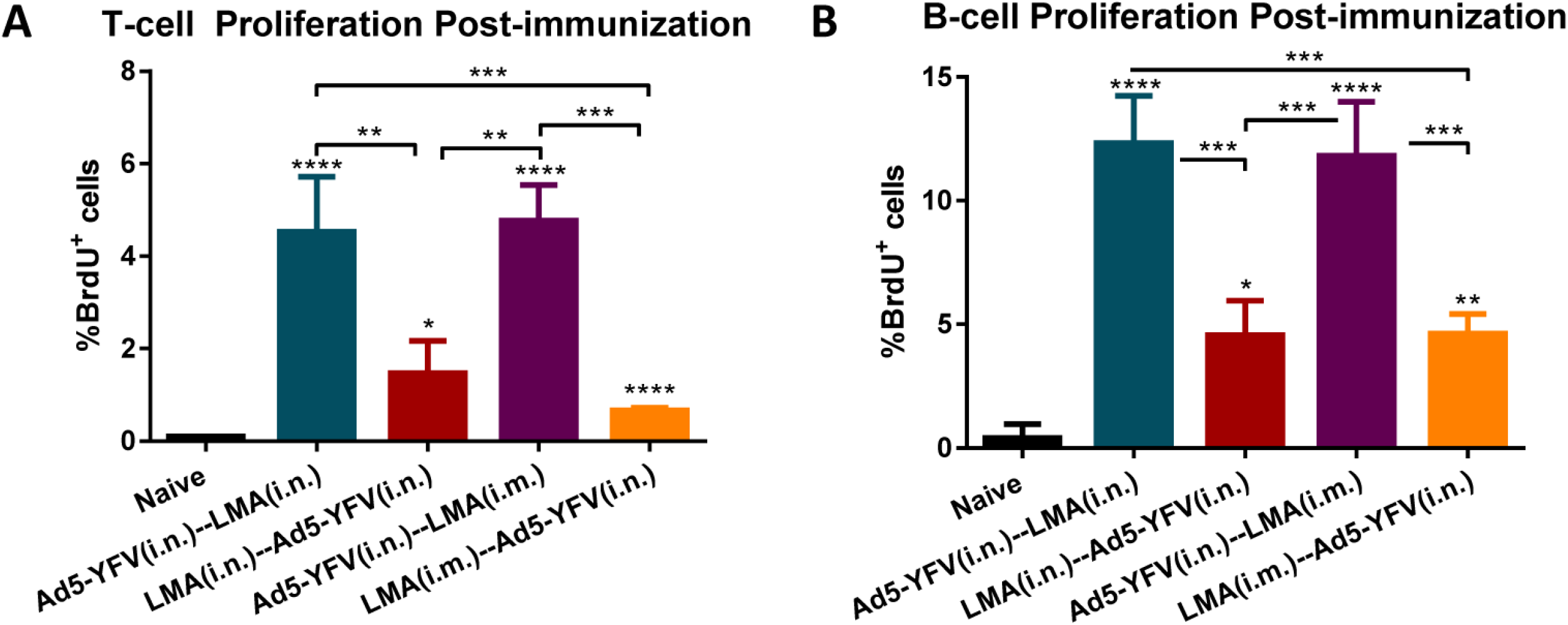
T- and B-cell proliferation in response to heterologous prime-boost vaccination during long-term study. Spleens were collected 21 days after the last vaccination dose from a cohort (n=5 per group) of immunized and naïve control mice as described in Fig. 5. The isolated splenocytes were stimulated with rF1-V (100 µg/ml) for 72 h 37° C and then BrdU was added at a final concentration of 10 µM during the last 18 h of incubation with rF1-V to be incorporated into newly synthesized DNA of the splenocytes. Subsequently, the BrdU-labeled splenocytes were surface stained for T- and B-cell markers followed by BrdU and 7-AAD staining. The splenocytes were then subjected to flow cytometry, and the percent of BrdU positive cells in CD3 (**A**) and CD19 (**B**) positive populations were calculated using FACSDiva software. Statistical significance was determined using One-way ANOVA with Tukey’s post-hoc test. Asterisks above columns represented comparison to the control group while asterisks with comparison bars denoted significance between the indicated groups. *p<0.05, **p<0.01, ***p<0.001, **** p<0.0001. Two biological replicates were performed, and data plotted. *In vitro* studies had 3 replicates.

### Conventional mice vaccinated with the 2-dose regimen were fully protected from CO92 and CAF^-^ challenges during a long-term study

The immunized mice along with naïve controls were then i.n. challenged with 100 LD_50_ of either CO92 or its CAF^-^ strain. As expected, all mice in vaccinated groups survived (with no clinical symptoms of the disease), while 100% of naïve control animals succumbed to infection (**Fig. 7A and B**, with up to 20% body weight loss). In addition, high plague bacilli (10^9^-10^10^ cfu/organ) were detected in various organs of diseased naïve control mice. In contrast, the inoculated *Y. pestis* was completely cleared from infected lungs of all the vaccinated mice after 28 days of the challenge (**Fig. 7C**). We have previously shown that mice vaccinated with either 2 doses of LMA or Ad5-YFV cleared the invading pathogen within 3 days p.i. (26, 27). At the end of the study (day 28 post-challenge), we collected BALF from all surviving mice, and all vaccinated and challenged animals had a significant level of F1-V specific IgA compared to PBS controls. (**Fig. 7D**).

**Figure 7:**
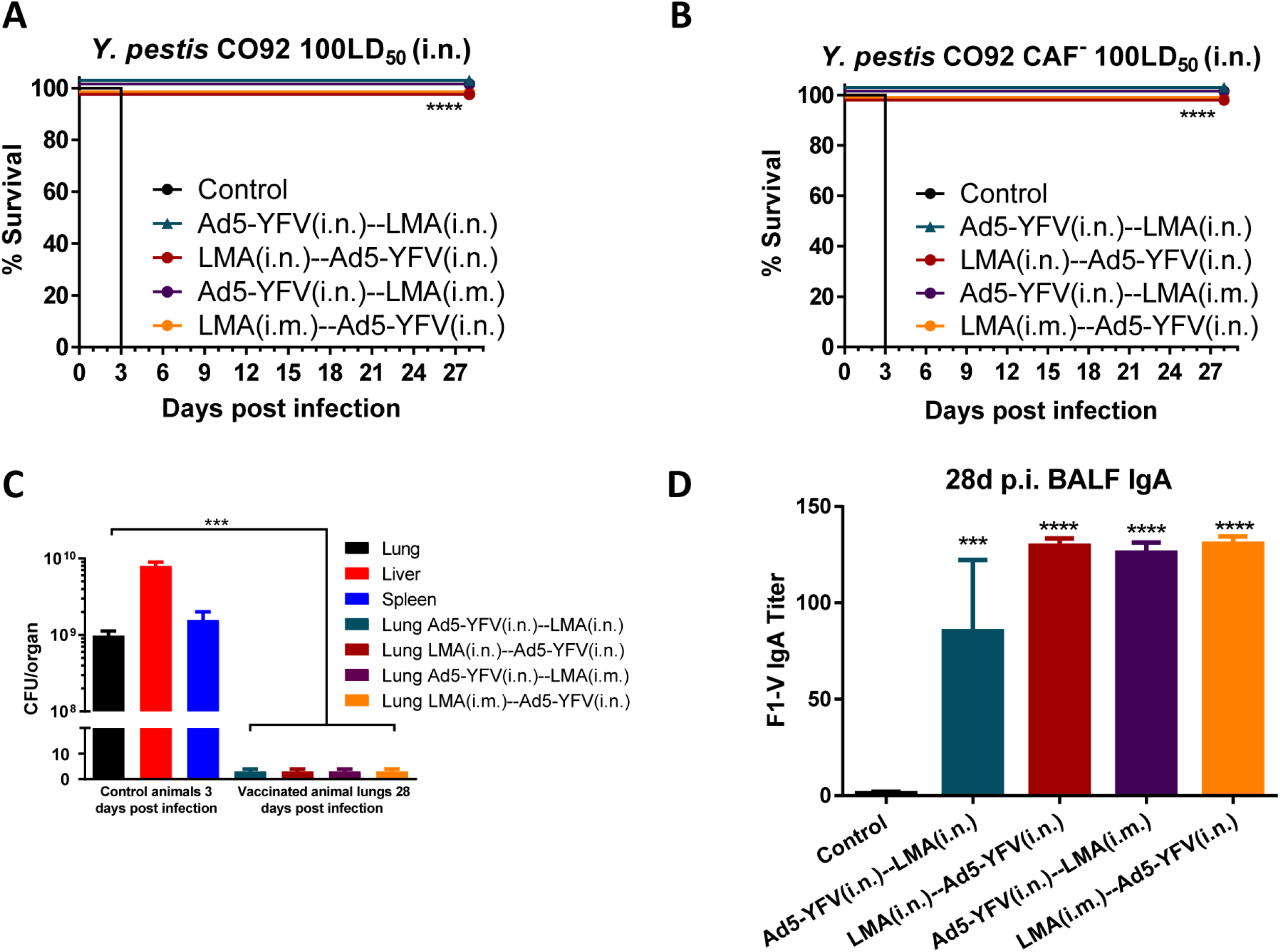
Heterologous prime-boost vaccinations provide protection to immunized mice in long-term study. A cohort (n=5-8 per group) of immunized and naïve control mice as described in Fig. 5. were challenged on day 105 of the study (84 days after 2^nd^ vaccination) with 100 LD_50_ of either CO92 (**A**) or its CAF^-^ strain (**B**) and monitored for morbidity and mortality for 28 days. On day 3 p.i., lungs, liver, and spleen were excised from all moribund mice to quantify bacterial load (**C**). At the end of the study (on day 28), BALFs and lungs were collected from the terminated animals. Lung homogenates were plated to determine the clearance of *Y. pestis* from the surviving animals (**C**). The collected BALFs were evaluated for IgA titers specific to F1-V by ELISA, and PBS-injected mice were used as controls (**D**). One-way ANOVA was used to determine significance between groups for bacterial burdens and antibody titers. While, Kaplan-Meier analysis with log-rank (Mantel-Cox) test was used for analysis of animal survivals. Asterisks represented the statistical significance compared to the control group or between the two groups indicated by a line. ***p<0.001, ****p<0.0001. Two biological replicates were performed, and data plotted. *In vitro* studies had 3 replicates.

We then examined sera of mice for IgA post vaccination and post challenge with CO92, and no differences in titers were noted (**Fig. S2**), suggesting infection did not further enhance serum IgA levels. A similar study will be performed in the future assessing IgA levels in BALF post vaccination and post challenge. Mice from the Ad5-YFV (i.n.)-LMA (i.n.) vaccinated group had slightly lower level of IgA than the other heterologous prime-boost vaccinated groups of animals, although this differences were not significant (**Fig. 7D**). Although it is expected that in pneumonic plague, IgA would be significantly contributing to host protection, some studies indicated minimal protective role of IgA (31, 45). However, whether the immune status of the host could be a contributing factor in IgA-associated protection is unclear and needs further investigation.

To gauge cell-mediated immunity of vaccinated mice in response to CO92 challenge, on day 3 p.i., splenocytes were isolated from both immunized and control animals and subjected to flow analysis after PMA/Ionomycin stimulation. We noted statistically higher IFNγ producing CD4^+^ and CD8^+^ T-cell populations across all of the vaccinated mice as compared to that of naïve control animals (**Fig. 8A and B**), and a similar trend was observed for the IL-17 producing CD4^+^ T-cells (**Fig. 8C**).

**Figure 8:**
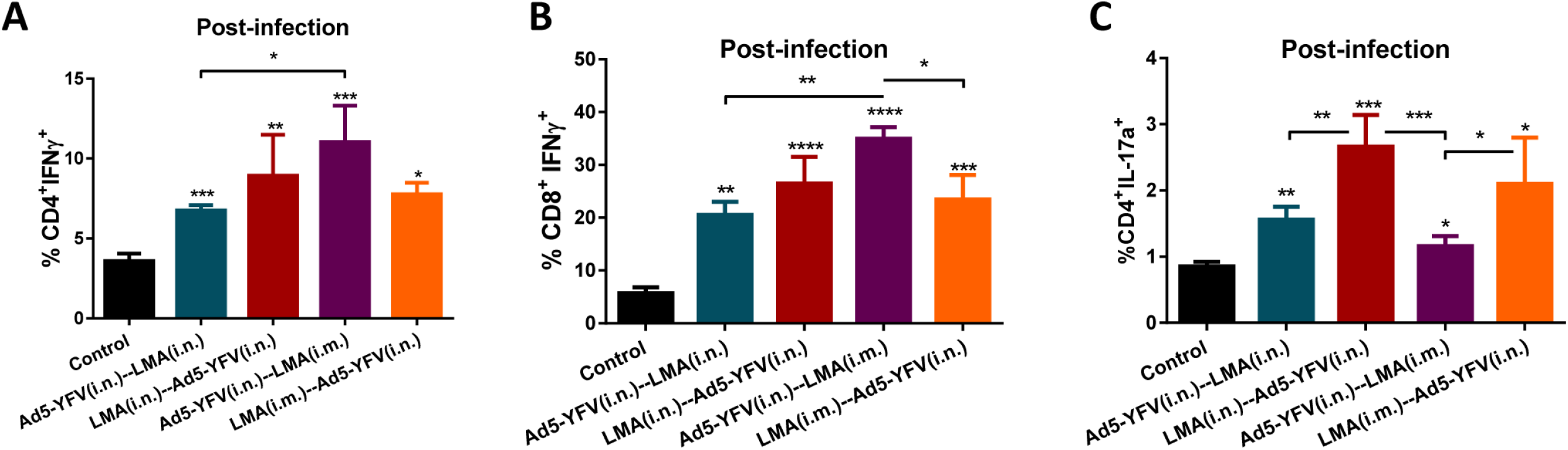
T-cell responses to CO92 challenge during long-term heterologous prime-boost vaccination study. A cohort (n=5 per group) of immunized and naïve control mice as described in Fig. 5. were challenged on day 105 of the study (84 days after 2^nd^ vaccination) with 100 LD_50_ of CO92. Spleens were harvested on day 3 p.i., and the isolated splenocytes were then stimulated with PMA, Ionomycin, and Brefeldin A. Cells were stained with T-cell surface markers CD3, CD4, and CD8 followed by intracellular IFNγ and IL-17A staining. Percentages of CD4^+^ IFNγ^+^ (**A**) and CD8^+^ IFNγ^+^ (**B**) and CD4^+^ IL-17^+^ cells (**C**) were shown. Cells were then analyzed by flow cytometry. Statistical significance was determined using One-way ANOVA with Tukey’s post-hoc test as well as Student’s t-test. Asterisks above columns represented comparison to the control group while asterisks with comparison bars denoted significance between other indicated groups. *p<0.05, **p<0.01, ***p<0.001, **** p<0.0001. Two biological replicates were performed, and data plotted. *In vitro* studies had 3 replicates.

More specifically, a significantly higher level of CD4^+^ IFNγ^+^ population was observed for mice immunized (purple bar) with Ad5-YFV (i.n.)-LMA (i.m.) in comparison to mice vaccinated (teal bar) with Ad5-YFV (i.n.)-LMA (i.n.) (**Fig. 8A**). On the other hand, a significant difference in CD8^+^ IFNγ^+^ population was only noticed between Ad5-YFV (i.n.)-LMA (i.m.) immunized group (purple bar) when compared to Ad5-YFV (i.n.)-LMA (i.n.) (teal bar) or LMA (i.m.)-Ad5-YFV (i.n.) vaccinated groups of mice (crimson bar) (**Fig. 8B**).

In terms of CD4^+^ IL-17^+^ population, mice immunized first with LMA either by the i.n. or the i.m. route generally revealed better levels than mice immunized with Ad5-YFV vaccine first. Significant differences were observed between LMA (i.n.)-Ad5-YFV (i.n.) immunized group (crimson bar) of mice when compared to groups vaccinated with either Ad5-YFV (i.n.)-LMA (i.n.) (teal bar) or Ad5-YFV (i.n.)-LMA (i.m.) (purple bar) as well as between groups immunized with LMA (i.m.)-Ad5-YFV (i.n.) (orange bar) and the group vaccinated with Ad5-YFV (i.n.)-LMA (i.m.) (purple bar) (**Fig. 8C**).

### Characterization of mice splenic cytokine and chemokine profiles in response to vaccination and CO92 challenge

To further evaluate cell-mediated immunity of mice in response to vaccination and infection, splenocytes collected from either post-immunization or post-challenge (**Fig. 5B**) were stimulated with rF1-V to examine cytokine/chemokine production. We divided up the cytokine/chemokine analysis into 3 panels: proinflammatory/anti-proinflammatory cytokines, Th1/Th2/Th17 associated cytokines, and chemokines. At post-vaccination time point, the overall proinflammatory cytokines in immunized mice were either slightly elevated (IL-6) or remained at the comparable levels (IL-1α and IL-1β) to those of naïve control mice (**Fig. 9**). However, in response to *Y. pestis* infection, significantly increased proinflammatory cytokines were observed only in naïve control mice, while the levels of these cytokines were low and almost unchanged in vaccinated animals (**Fig. 9**). For the anti-proinflammatory cytokine IL-10, a significant increase was generally observed in all of the immunized mice as compared to that of naïve controls at the post-immunization time point. In response to *Y. pestis* infection, the level of IL-10 was significantly elevated in both naïve control mice and in animals immunized first with LMA (crimson and orange bars); however, it remained unchanged in mice immunized first with Ad5-YFV (teal and purple bars), and was significantly lower than that of naïve control at the post challenge time point (**Fig. 9**).

**Figure 9:**
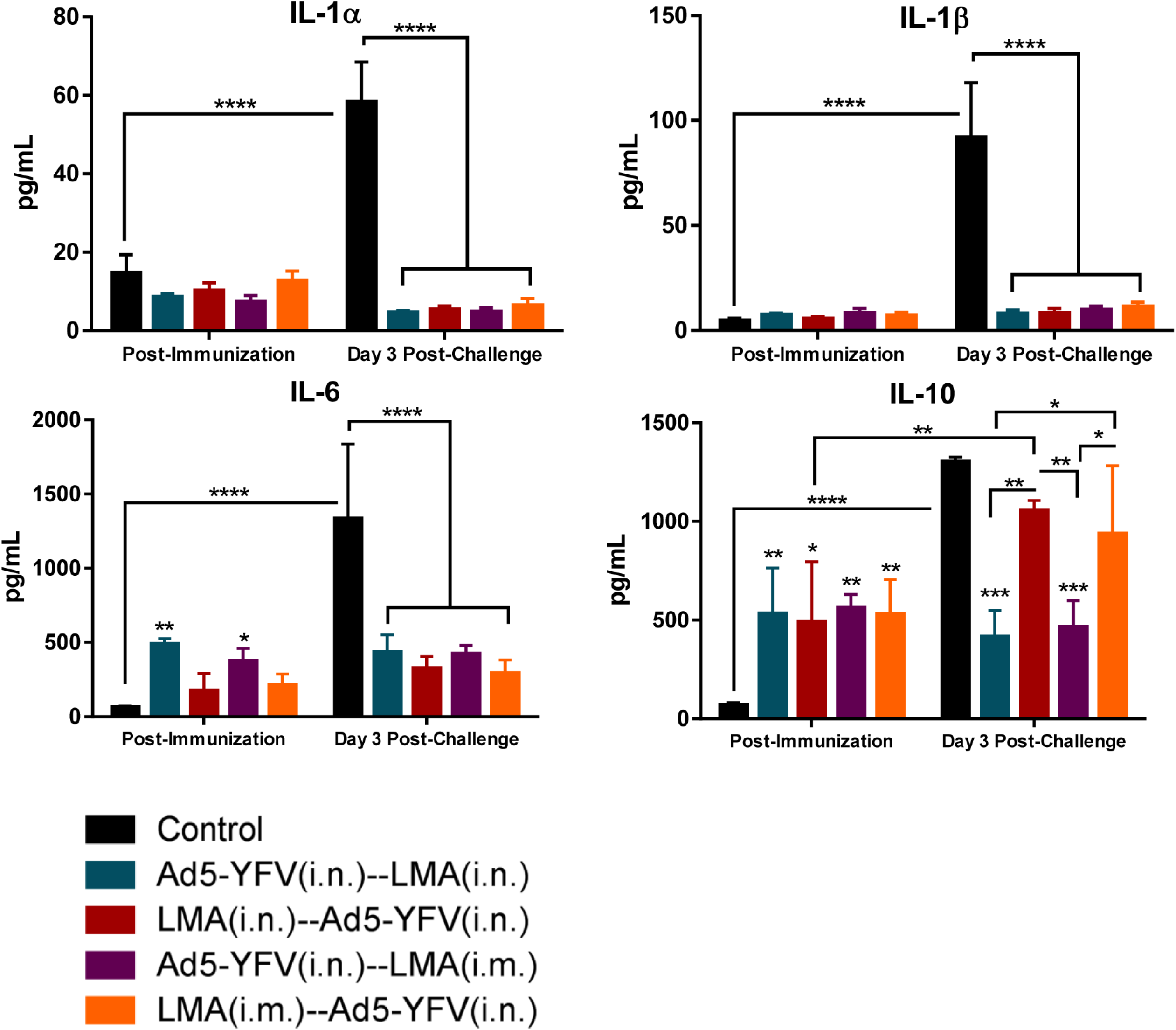
Splenocyte proinflammatory and anti-proinflammatory responses during long-term heterologous prime-boost vaccination study. The splenocytes isolated from mice described in Fig. 6 (post immunization) and Fig. 8 (post CO92 challenge) were further stimulated with rF1-V (100 µg/mL) for 3 days. The cytokines in the culture supernatants were analyzed by using Bioplex-23 assay and expressed as the arithmetic means ± standard deviations. The proinflammatory and anti-proinflammatory cytokines are shown. Statistical significance was determined using Two-way ANOVA with Tukey’s *post hoc* test to compare multiple time-points or student t-test to compare 2 groups within the same time-point. Asterisks above columns represented comparison to the control group, while horizontal bars represented differences between test groups. *p<0.05, **p<0.01, ***p<0.001, **** p<0.0001. Two biological replicates were performed, and data plotted. *In vitro* studies had 3 replicates.

In contrast to proinflammatory cytokines, the Th1- and Th2-related cytokines such as IL- 2, IL-12(p70), IFNγ, IL-4, IL-5, and IL-13 were generally increased in all immunized mice as compared to that of naïve controls at the post-immunization time point (**Fig. 10**). On the other hand, IL-2, IL-12(p70), IFN-γ and IL-4 levels were sustained at higher levels in all immunized groups in response to CO92 challenge. The levels of IL-5 and IL-13 were only maintained higher in mice first immunized with LMA (crimson and orange bars) but subsided to the level of naïve controls in mice first vaccinated with Ad5-YFV (teal and purple bars) at the post-challenge time point (**Fig. 10**). The decline of Th2 cytokines IL-5 and IL-13 in mice immunized first with Ad5-YFV at post-challenge time point might reflect its Th1 bias as also shown above in analysis of IgG isotyping (**Fig. 4B and Fig. 5D**).

**Figure 10:**
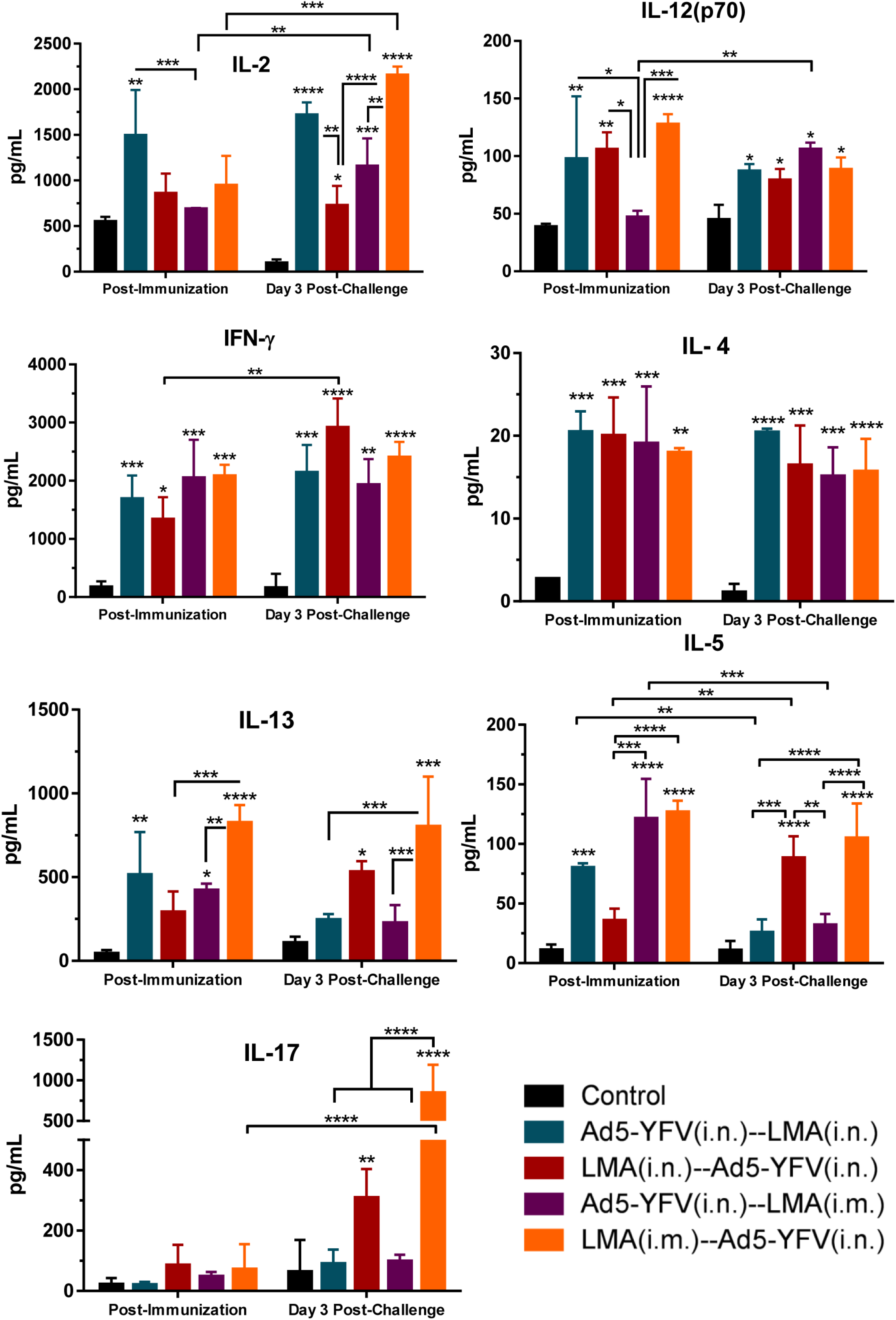
Splenocyte Th1/Th2/Th17 cytokine responses during long-term heterologous prime-boost vaccination study. Cytokine analysis was performed as described in Figure 9. Th1/Th2/Th17 cytokine responses are shown. Statistical significance was determined using Two-way ANOVA with Tukey’s *post hoc* test to compare multiple time-points or student t-test to compare 2 groups within the same time-point. Asterisks above columns represented comparison to the control group, while horizontal bars represented differences between test groups. *p<0.05, **p<0.01, ***p<0.001, **** p<0.0001. Two biological replicates were performed, and data plotted. *In vitro* studies had 3 replicates.

For Th17 response, slight increases in the levels of IL-17 were only observed in mice immunized first with LMA (crimson and orange bars) as compared to that of the naïve controls at post-immunization. In addition, the IL-17 level in these mice was further elevated in response to *Y. pestis* challenge. However, the IL-17 levels were consistently maintained at a low level and was comparable with that of naive controls at both time points (post immunization and post challenge) in mice vaccinated first with Ad5-YFV (teal and purple bars) (**Fig. 10**). It should be mentioned that it is difficult to precisely correlate percentage of T cells positive for IL-17 as assessed by flow cytometry and IL-17 secreted by these cells based on Bioplex as the cells were stimulated with different agents, namely PMA or F1-V; however, a corelative trend should be expected as shown in **Figs. 8C and 10**.

In analysis of chemokine production, the overall level of chemokines in all immunized mice were low and comparable with their corresponding naïve controls at the post immunization time point except the GM-CSF which was significantly elevated (**Fig. 11**). In response to *Y. pestis* infection, the CXCL1, RANTES, and G-CSF behaved similarly to proinflammatory chemokines and were significantly increased only in naïve control mice but were at low levels and largely unchanged in all immunized and challenged animals (**Fig. 11**). In contrast, CCL2, CCL4, and GM-CSF had generally elevated levels in all immunized mice as compared to that of naïve control at post-challenge time point. The exception being CCL4 in mice immunized with LMA i.m. (purple and orange bars) which were at a similar level to that of naïve control challenged mice (**Fig. 11**).

**Figure 11:**
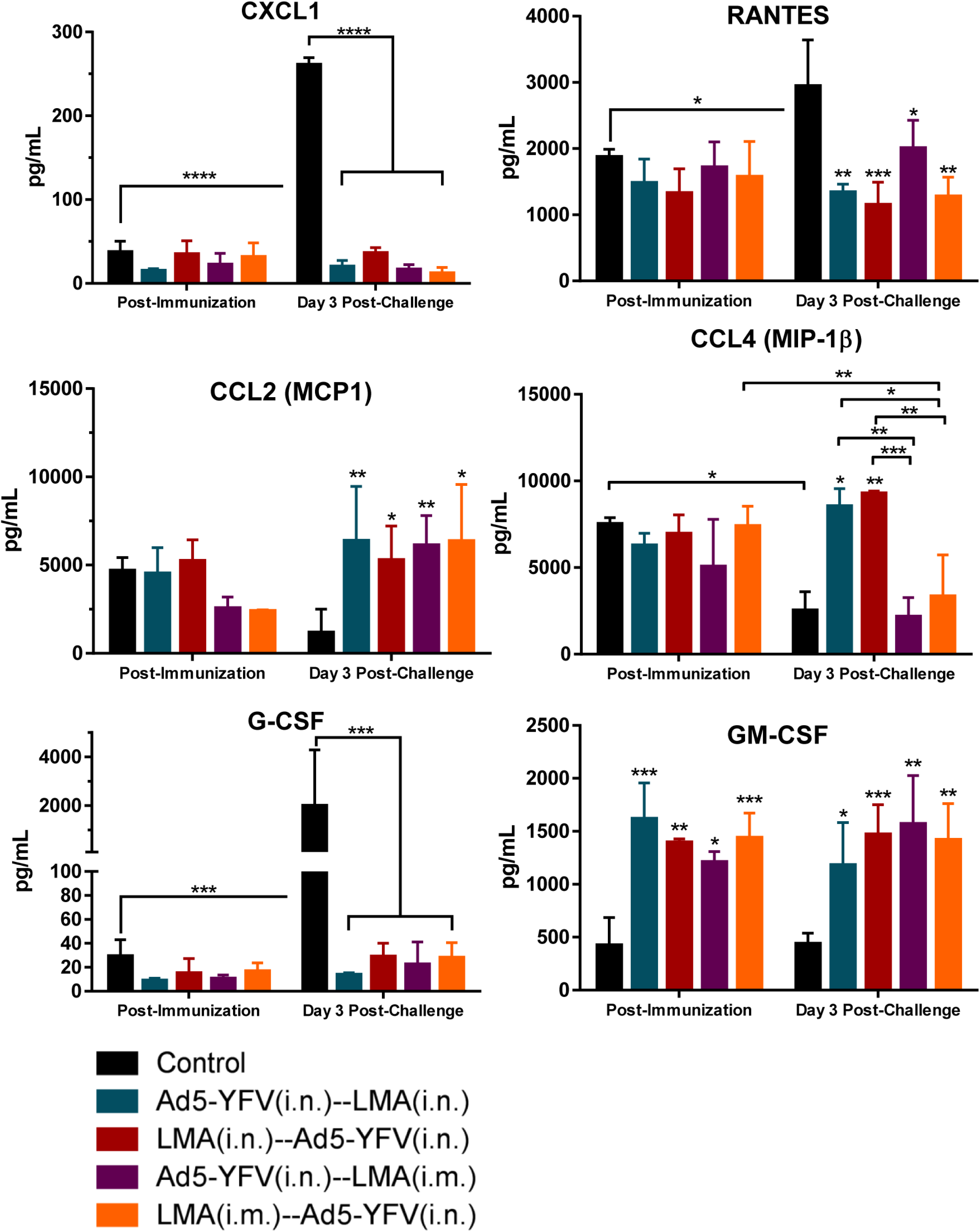
Splenocyte chemokine responses during long-term heterologous prime-boost vaccination study. Chemokine analysis was performed as described in Figure 9. Chemokine responses are shown. Statistical significance was determined using Two-way ANOVA with Tukey’s *post hoc* test to compare multiple time-points or student t-test to compare 2 groups within the same time-point. Asterisks above columns represented comparison to the control group, while horizontal bars represented differences between test groups. *p<0.05, **p<0.01, ***p<0.001, **** p<0.0001. Two biological replicates were performed, and data plotted. *In vitro* studies had 3 replicates.

## Discussion

After the 2017 outbreak of plague in Madagascar, the WHO released a target product profile that outlined the desired characteristics for a potential plague vaccine (46). These characteristics included: at most 2-dose vaccination schedule, long-lasting protection with humoral and cell-mediated responses, possibility of a needle-free administration, and a robust safety profile including potential use in pregnant women, children, and immuno-compromised individuals (46).

We recently have developed two plague vaccine candidates: a live-attenuated vaccine LMA and an Ad5 viral vector-based vaccine Ad5-YFV; individually both of them provided complete protection to immunized animals against challenge with CO92 (25–28). In response to the WHO requirements, here, we first further investigated the safety of the LMA vaccine in an iron-overload condition and then in Rag1 KO mice (47). In live-attenuated plague strains that rely on pigmentation locus mutations such as KIM/D27 and EV76, hereditary hemochromatosis has been shown to restore bacterial virulence (48). While the LMA mutant has intact pigmentation locus with functional T3- and T6-secretion systems (19) (data not shown).

Unlike KIM/D27 strain which exhibited more virulence under an iron-overload environment, LMA vaccine was not affected by the presence of more iron in mice (Fig. 1). The *Rag1* gene defect in humans is associated with a broad spectrum of clinical and immunological phenotypes, and is one of the major causes of human immune deficiency (PID) (49, 50). The Rag1 KO mice receiving up to 2.0 x 10^6^ CFU of LMA, which is equivalent to 20,000 LD_50_ of CO92, did not exhibit any clinical symptoms of the disease and the LMA mutant rapidly cleared from these mice (Fig. 2). These data demonstrated a high degree of attenuation imparted by the selected mutations in LMA and provided indication that the vaccine would be suitable for use in immunocompromised individuals.

We then implemented a heterologous immunization strategy in which both LMA and Ad5-YFV vaccines were delivered either simultaneously (1-dose regimen) or in a prime-boost format (2-dose regimen), as we hypothesized such a strategy would have several advantages. First, the use of two rationally-designed vaccines, which are based on different principles, would complement each other from their own potential disadvantages such as limited plague antigens in the Ad5-YFV vaccine versus the LMA vaccine which would provide immune responses to plethora of antigens. Second, the use of an Ad5-YFV vaccine as the first dose would negate any safety concerns of employing LMA as the second booster dose. Third, the heterologous immunization scheme is expected to mount unique and durable immune responses by integrating characteristics of each of the two vaccines, and thus, leading to superior protection in a broader human population. Fourth, the 1-dose regimen would shorten the immunization course and considered ideal to be used in emergency situations such as a plague outbreak or during a bioterrorist attack.

Indeed, all our heterologous immunizations (irrespective of the order of vaccine delivery and the routes of administration) induced robust immune responses in mice and provided full protection to animals against the lethal challenges of both CO92 and its CAF^-^ mutant. The exception was the group of mice that received Ad5-YFV and LMA vaccines simultaneously by the i.n. route and had 50 to 63% survival rates during CO92 or its CAF^-^ strain challenges (Fig. 3). This relatively lesser protection rate in mice was correlated well with lower F1-V specific antibody titers in mouse serum as compared to animals simultaneously immunized with Ad5-YFV and LMA *via* different routes, *i.e.*, i.n. and i.m., respectively (Fig. 4A). Further, there could be interference in triggering protective immune responses when both LMA and Ad5-YFV vaccines were delivered simultaneously in the lungs. It is also plausible that a stronger innate immunity developed in the lungs due to administration of two vaccines simultaneously, resulted in their respective rapid clearance, thus decreasing overall immunogenicity. Similar results were reported in a study in which tuberculosis vaccine BCG was used in combination with other TB subunit vaccines (*e.g*., 85A, E6, and TB10.4) in a heterologous format. Simultaneous administration of them *via* different routes (pulmonary and parental) induced both pulmonary and systemic immunities resulting in better protection compared to animals that were simultaneously vaccinated *via* the same route (51). Interestingly, the Ad5-YFV (i.n.) and LMA (i.m.) simultaneous vaccination combination also exhibited immune response characteristics of both Ad5-YFV (Th1) and LMA (Th17) vaccines (Fig. 4B and E) with complete protection of mice against challenges with CO92 and its CAF^-^ mutant (Fig. 3E and F). Thus, 1-dose regimen of two vaccines provided us with an excellent tool to combat plague during emergency situations.

It is difficult to discern which ones of our 2-dose heterologous regimens were better as all of them provided full protection to the immunized animals with comparable levels of F1-V specific antibodies and an overall similar cytokine profiles. However, an intriguing phenomenon emerged in that the immune profile of 2-dose immunization was likely dictated by the vaccine which was administered first.

More specifically, when the Ad5-YFV vaccine was delivered first, Th1 immune response was more pronounced based on F1-V specific IgG2a/IgG1 antibody ratio (Fig. 4B) and the splenic cytokine profiles (Fig. 10). Likewise, when LMA vaccine was administered first followed by that of Ad5-YFV, the Th2 immune response was favored (Fig. 3B and Fig. 8C) along with that of Th17 (Fig. 8C and Fig. 10) response. Further, clear differences were noted in terms of T- and B-cell proliferation when mice were immunized first with the Ad5-YFV vaccine in response to stimulation with rF1-V and had higher LcrV antibody titers when compared to animals receiving LMA vaccine first during the long-term study (Fig. 5E and Fig. 6). Therefore, in this regard, delivering the Ad5-YFV vaccine first followed by that of LMA vaccine would be preferred, and is much more attractive based on safety and that both vaccines can be administered i.n., thus developing a more acceptable adjuvant- and needle-free administration protocol from the public health prospective.

The heterologous vaccination strategies have been previously used for diseases in which cell-mediated responses were particularly important for protection such as in patients with HIV and malaria (52). Recently, during the COVID-19 pandemic, the heterologous prime-boost vaccination has been emphasized mainly due to the shortage of available COVID-19 vaccines (53). The intentional design has also been reported in the Russian Sputnik vaccines which use two different adenovirus vectors (type 5 and type 26), and thus far, this is the only heterologous prime-boost vaccine to be licensed for human use (54–56). Further, with many COVID-19 vaccines under development based on different platforms and strategies, and the likely need for additional boosters due to the emerging COVID-19 variants, have led to more enthusiasm in investigating the advantages of heterologous prime-boost approach over homologous boost strategy.

Indeed, it was shown that using a heterologous boost of either adenoviral-vectored vaccine or mRNA-based COVID-19 vaccines improved neutralizing antibody titers with induction of stronger Th1 responses than a homologous boost of inactivated SARS-CoV-2 vaccines (57). Similarly, using a mRNA based COVID-19 vaccine as a booster (BNT162b2) in a heterologous approach instead of using a homologous adenoviral booster (ChAdOx1-nCov-19) resulted in increased neutralizing antibody titers and SARS-CoV-2 specific T-cells (58). Although there is no data for direct comparison between heterologous and homologous immunization approach with LMA and Ad5-YFV vaccines; we did notice that the antibody titers to the individual antigen LcrV was the lowest among three tested antigens (LcrV, F1, and YscF) in mice immunized solely with the Ad5-YFV vaccine (26). In contrast, when mice were immunized with both LMA and Ad5-YFV vaccines in the heterologous format, a comparable level of antibodies to all three antigens was observed (Fig. 5E).

Recently, a prime and pull immunization regimen has been investigated in a variety of vaccines with success (59–61), which implicates that the immune response at a mucosal site can be triggered by the administration of an antigen to a distant mucosal site. In this strategy, a parenteral vaccination raises systemic cellular response (prime) followed by a mucosal delivery of chemokine or vaccine that directs the tissue targeting of the prime-activated circulating T cells (pull) (62). In our study, mice i.m. immunized with LMA followed by i.n. delivery of Ad5-YFV vaccine exactly fits this category and should be further investigated. Although significant increases in splenic T cell population (both CD4 and CD8) was observed in all immunized mice as compared to the naïve controls, no significant differences among all of the immunized groups of mice was observed. In addition to the circulating T cells, the hallmark of prime-pull immunization is the elevated local T cell population, especially the resident memory T cells (TRM), at the mucosal sites (59–61). TRM occupies tissues without recirculating and provides a first response against infections and accelerates pathogen clearance (63). Therefore, it is possible that the local level (lungs) of T cells varies among our different heterologous immunized groups of animals and needs to be further studies.

A new vaccination strategy by combination of both simultaneous and prime-boost immunizations has been used for H1N1 signal minus influenza vaccine (S-FLU) in a pig model (64). During the prime-boost immunization course, the study has shown i.m. only immunized pigs generated a high titer of neutralizing antibodies but poor T cell responses, whereas aerosol solely induced powerful respiratory tract T cell responses but a low titer of antibodies. However, immunization of animals with S-FLU *via* both i.m and aerosol routes simultaneously during the prime-boost immunization course generated high antibody titers and strong local T cell responses with the most complete suppression of virus shedding and the greatest improvement in pathology (64). This strategy has been highly recommended for TB vaccines as well (51). Considering the Ad5-YFV (i.n.) and LMA (i.m.) simultaneous combination was the only one among all immunized groups that showed the immune characteristics of both Ad5-YFV (Th1) and LMA (Th17) (Fig. 4B and E), it will be exciting to carry out a similar experiment with LMA and Ad5-YFV vaccines in lieu of S-FLU in the future.

Finally, cytokine/chemokine production (*e.g.,* IL-1α, IL-1β, and IL-6) from splenocytes of unvaccinated mice 72 h p.i. with CO92 indicated a highly inflammatory environment with neutrophil chemoattractants (CXCL1, RANTES, and G-CSF) at elevated level (Fig. 11). None of these cytokines/chemokines were elevated in any of the combinations of heterologous prime-boost vaccinated mice, indicating inability of *Y. pestis* to cause immune dysregulation during early stages of infection. Conversely, immunized and infected mice in all heterologous prime-boost groups had higher levels of CCL2, and some groups had higher CCL4, which could be essential in recruiting monocytes to clear the invading the pathogen. Importantly, mice immunized with the LMA vaccine first had the highest amount of secreted IL-17 during the post-infection time point, which combined with decrease in CXCL1, RANTES, G-CSF, and increases in CCL2, CCL4, and GM-CSF could have a critical role during productive versus non-productive stages of pneumonic plague.

In general, by using heterologous vaccination strategy, we have clearly demonstrated that the combination of vaccines, the route and timing of administration, as well as the length of the vaccination schedule are all crucial factors that affect efficiency and safety of vaccinations. It is important to reiterate that almost all our heterologous combinations offered complete protection from high-dose challenges of both CO92 and its CAF^-^ strain, and each one of them has its own characteristics that can fit for different scenarios. Importantly, the 2-dose regimens especially the Ad5-YFV (i.n.) and LMA (i.n.) combination is ideal for the routine immunization in plague endemic regions, while the simultaneous approach with Ad5-YFV (i.n.) and LMA (i.m.) would be beneficial for vaccination in response to emergency situations.

Our future studies will address three important questions and include: **1)** comprehensively assessing potency of memory responses (T- and B-cells) that would navigate us on dosing strategies; **2)** assessing antibody potency in neutralizing *Y. pestis* infection, which would also support our heterologous prime-boost approach with two vaccines and potentially could reveal other additional differences that are important in humans; and **3)** testing heterologous prime-boost strategy in humanized mouse model and non-human primates to gauge superiority of our approach compared to competing vaccine candidates, and that our Ad5-YFV and LMA combination would be highly efficacious in humans.

## Methods

### Bacterial strains and vaccines

A fully virulent human pneumonic plague isolate, the parental *Y. pestis* strain (CO92), was obtained from BEI Resources (Manassas, VA). The F1-negative strain (CAF^-^) was created in our laboratory by deletion of partial *caf1A* and most of the *caf1* gene from CO92. The mutant strain retained its virulence in both pneumonic and bubonic plague animal models (39). The live-attenuated vaccine candidate (LMA) is a triple deletion mutant of CO92 in which genes encoding Lpp, MsbB, and Ail, were deleted (19). The KIM/D27 strain of *Y. pestis* deleted for the pigmentation locus required for iron acquisition from the host was used in an iron-overload experiment performed in mice (65). The Ad5-YFV is an human replication-defective adenovirus type 5 vector-based vaccine containing genes for three plague antigens: F1, LcrV, and YscF (25). All studies involving *Y. pestis* were performed in Tier 1 select agent laboratories at UTMB in the Galveston National Laboratory (GNL), Galveston, TX.

A large batch of Ad5-YFV vaccine [1×10^16^ virus particles (v.p.)/batch, aliquoted in 1 ml at 1×10^12^ v.p. and stored at −80°C] was prepared from a 20-liter suspension culture of HEK293 cells in a chemically defined, protein-free CD-293 medium. The vaccine was purified at the Baylor College of Medicine Vector Development Laboratory and by our company partner in collaboration with Lonza, Houston, TX, under good laboratory practice conditions. This batch of vaccine was used for our subsequent studies in mice and non-human primates (25, 26). Likewise, a large batch of the LMA vaccine (2-liters) was prepared under a highly regulated quality control system in GNL biosafety level 3 (BSL-3) suite, by growing in Heart Infusion Broth (HIB) overnight at 28°C as a shake flask culture (180 rpm). The culture was centrifuged, washed with HIB, and resuspended to 1/20^th^ the original volume. The culture was aliquoted (500 µl, ∼1×10^9^ colony forming units [CFU]/ml) with 25% glycerol and stored at −80°C. Titers of the vaccines were confirmed before and after each inoculation, and the same batches of the vaccines were used throughout our earlier and these studies (19, 27).

### Animals

Outbred Swiss-Webster (female) and inbred C57BL6 Rag1 Knockout (KO, lacking mature T- and B-cells, male and female)) mice (6-8 weeks) were purchased from Jackson Laboratory (Bar Harbor ME). All experiments were conducted in the animal biosafety level 3 (ABSL-3) facility at UTMB in the GNL. Studies were ethically performed under an approved Institutional Animal Care and Use Committee protocol.

### Iron-overload studies

Swiss-Webster mice (n=5/group) were injected with 75 µg of ferrous chloride (FeCl_2_·4H_2_O, Sigma-Aldrich Inc., St. Louis, MO) by the intraperitoneal (i.p.) route (65) and then challenged i.n. with either the KIM/D27 strain or the LMA vaccine strain (2×10^5^-5×10^6^ CFU/50 µl). A similar number of untreated mice (without FeCl_2_·4H_2_O) were also infected and served as controls. The animals were observed for body weight loss, other clinical signs of the disease (ruffled fur, hunched back, lethargy, lack of grooming, sunken eyes, squinting of eyes with ocular and or nasal discharge, open-mouth breathing, gasping for air), and morbidity over a period of 14 days.

### Rag1 KO mice studies

Rag1 KO mice (n=10, males or females) were infected with 4 LD_50_ of CO92 by either i.n. or i.m. routes and observed for morbidity and mortality. For inbred mice, 1 LD_50_=10 or 100 colony-forming units (CFU) when delivered by the i.m. or the i.n. route, respectively. Lungs, liver, and spleen were excised from moribund mice and homogenized in 1-2 mL of phosphate-buffered saline (PBS). Homogenates were 10-fold serially diluted and plated on sheep blood agar (SBA) plates to quantify bacterial load. For the LMA vaccine studies, Rag1 KO mice (10 for each infection route) were infected with 2.0 x 10^6^ CFU by either the i.n. or the i.m. route. A cohort of 10 mice (5 from each infection route) was sacrificed on day 5 post-infection (p.i.), and bacterial loads in spleen and at the initial infection sites (lungs or the muscle) were determined as described above. The remaining 10 mice were observed for signs of disease for 28 days, and the survivors were then i.n. challenged with 6 LD_50_ of CO92. On day 3 post-challenge, mice were bled to collect serum, and lungs from 5 moribund animals were excised on day 4 to quantify bacterial load.

### Heterologous vaccination and challenge studies

Swiss-Webster mice were immunized with Ad5-YFV and LMA vaccines in either a 1- or 2-dose regimens. In a 1-dose regimen, both Ad5-YFV and LMA were delivered simultaneously, while in a 2-dose regimen, Ad5-YFV and LMA vaccines were administered 21 days apart in various order and route combinations. Mice receiving PBS were used as controls. The vaccination doses were 1.2 x 10^10^ plaque forming units (PFU)/40 µL for Ad5-YFV and 2.0 x 10^6^ CFU/50 µL for the LMA vaccine (26, 27).

For short-term studies, the immunized and naïve control mice (n=8-10) were retro-orbitally bled on day 21 after the last dose of immunization. The animals were then i.n. challenged with 100 LD_50_ of CO92 or CAF^-^ mutant on day 24 after completion of the vaccination course. For Swiss-Webster mice, 1 LD_50_=500 CFU of CO92 or CAF^-^ by the i.n. route (39, 66). In a separate experiment, mice (n=5) were similarly immunized with various combinations as described above; however, the group in which LMA and Ad5-YFV vaccines were administered simultaneously *via* the i.n. route was excluded because of interference in generating immune responses and reduced animal protection after bacterial challenge (see results). On day 21 after completion of the immunization course, mice were euthanized, and spleens collected for the evaluation of cell-mediated immunity.

For long-term studies, mice were immunized with the 2-dose regimen only, and animals receiving PBS were used as controls. Blood and spleens were collected from 5 mice in each immunized and control group on day 63 (42 days after last immunization dose), and the rest of mice were challenged with 100 LD_50_ of either CO92 or its CAF^-^ strain on day 105 (84 days after last immunization dose). On day 3 post challenge with CO92, organs (spleen, lungs, and liver) were collected from all moribund animals to quantify bacterial load, and spleens were also excised from 5 alive animals (after euthanization) of each group. At the end of the experiment, bronchoalveolar lavage fluid (BALF), spleen and lungs or muscle were collected from all the surviving animals. The isolated sera and BALFs were filtered using 0.1 µm filter cartridges (Millipore Sigma Life Science Center, Burlington, MA) and sterility confirmed before performing subsequent experiments at a lower biocontainment level. The collected organs were either used for quantitation of the bacterial load or for the evaluation of cell-mediated immunity.

### Antibody titer analysis

Antibody titers were measured by performing indirect Enzyme-linked immunosorbent assay (ELISA) (26). Briefly, MaxiSorp ELISA plates (NUNC, Rochester, NY) were coated with 100 ng of recombinant fusion protein (rF1-V) (BEI Resources) or individual plague antigens rF1, rLcrV, or rYscF in carbonate buffer (100 µL) at 4°C overnight. Non-coated antigens were removed with 3 washes of Dulbecco’s PBS (DPBS) with 0.05% Tween 20. Plates were then blocked with 1% powdered milk (EMD Chemicals Inc., Gibbstown, NJ) in DPBS for 1 h at room temperature. After 3 more washes, sera or BALFs were 2-fold serially diluted and incubated for 1-h at room-temperature. Plates were again washed 3 times and then horseradish peroxidase (HRP)-conjugated secondary anti-mouse antibodies for IgG, IgG1, IgG2a or IgA (Southern Biotech, Birmingham, AL), diluted at 1:8000, were added and incubated for 1 h at room temperature. Plates were washed 3 times and then 100 µL of TMB (3,3’,5,5’- Tetramethylbenzidine) substrate was added for 5-15 min at room temperature. Colorimetric reaction development was stopped using 2N H_2_SO_4_. Absorbance was then measured at 450 nm using a Versamax tunable microplate reader (Molecular Devices San Jose, CA).

### T-cell phenotypes

Spleens collected from both immunized and control mice were smashed and passed through a 70 µm cell strainer to obtain single cell suspension in RPMI 1640 cell culture medium. Splenocytes were then seeded into 24 well tissue culture plates at a density of 2.0 x 10^6^ cells/well. Four wells/mouse/plate were treated with ionomycin (750 ng/mL, calcium ionophore), PMA (phorbol 12-myristate 13-acetate, protein kinase C activator, 50 ng/mL), and Brefeldin A (5 µg/mL) for 5 h at 37° C in a 5% CO_2_ incubator. Stimulation of splenocytes with PMA and ionomycin leads to activation of several intracellular signaling pathways, bypassing the T cell membrane receptor complex, resulting in strong T cell activation and production of a variety of cytokines. Splenocytes were then blocked with anti-mouse CD16/32 antibodies (BioLegend, San Diego, CA) followed by staining with Fixable Viability Dye eFluor™ 506 (eBioscience, San Diego, CA) and APC anti-mouse CD3e (eBioscience), PE/Dazzle 594 anti-mouse CD4 (BioLegend), FITC anti-mouse CD8 (BioLegend) for CD3, CD4, and CD8 T-cell surface markers, respectively. Cells were then permeabilized for intracellular staining with PerCP/Cy5.5 anti-mouse interferon (IFN)-γ, PE/Cy7 anti-mouse IL-17A (BioLegend), and analyzed by flow cytometry.

### Cell proliferation and cytokine production

To measure T- and B-cell proliferation, bromodeoxyuridine (BrdU), a thymidine analog, incorporation method was used. Briefly, the isolated splenocytes were seeded in duplicate 24 well tissue culture plates (1.0 x 10^6^ cells/well) and stimulated with rF1-V fusion protein (100 µg/ml) for 72 h at 37°C. The BrdU (BD Bioscience, San Jose, CA) was then added into the wells of one of the plates at a final concentration of 10 µM during the last 18 h of incubation with rF1-V to be incorporated into newly synthesized DNA of the splenocytes (67, 68).

Subsequently, the BrdU-labeled splenocytes were surface stained for T-cell (CD3e-APC; eBioscience) and B-cell (CD19-eFluor450, ThermoFisher Scientific, Grand Island, NY) markers after blocking with anti-mouse CD16/32 antibodies (BioLegend). Cells were then permeabilized and treated with DNase to expose BrdU epitopes followed by anti-BrdU-FITC and 7-AAD (7-amino-actinomycin D) staining by using BD Pharmingen FITC BrdU Flow Kit (San Jose, CA). The splenocytes were then subjected to flow cytometry, and data analyzed as we previously described (28). The percent of BrdU positive cells in CD3 and CD19 positive populations were calculated using FACSDiva software.

To assess cytokine production, cell supernatants were collected from a duplicate plate above after stimulation with rF1-V (100 µg/ml) for 72 h at 37°C. For these studies, we used *Y. pestis* specific antigens for stimulation to confirm flow cytometry data. Cytokines in the supernatants were then measured by using Bio-Plex Pro Mouse Cytokine 23-plex Assay (Biorad Laboratories, Hercules, CA) following the manufacturer’s standard protocol.

### Statistical Analysis

One-way or Two-way analysis of variance (ANOVA) with Tukey’s *post hoc* test or the Student’s t-test was used for data analysis. We used Kaplan-Meier with log-rank (Mantel-Cox) test for animal studies, and P values of ≤0.05 were considered significant for all the statistical tests used. The number of animals per group is described in each figure or figure legend and two biological replicates were performed. All *in vitro* studies were performed in triplicates.

## Acknowledgements

P.B.K. was supported in part by Biodefense T32 Training Fellowship (T32-AI060549) awarded to A.K.C. We thank Dr. Yuejin Liang, Department of Microbiology and Immunology, in the analysis of the flow cytometry data. These studies were supported in part by funding from the NIH (AI153524 and AI071634) grants as well as UTMB Technology Commercialization Program and John S. Dunn Foundation funding awarded to A.K.C. Contributions of our company partner is greatly acknowledged.

---

**Figure S1. Parental *Y. pestis* CO92 causes clinical disease in Rag1 KO mice.** C57BL6 Rag1 KO mice (n=4-5 each route) were infected with 4 LD_50_ of CO92 by either (i.n.) or (i.m.) route and the mortality of animals recorded and plotted (**A**). Lungs, liver, and spleen were removed from the moribund animals to quantify the bacterial loads (**B**). Kaplan-Meier analysis with log-rank (Mantel-Cox) test was used for analysis of animal survival (**A**). Asterisks represent the statistical significance between two groups indicated by a line and *** p<0.001 (**B**). Two biological replicates were performed, and data plotted.

**Figure S2. No significant differences in F1-V specific serum IgA are observed in vaccinated mice before and after infection.** Mice were immunized with Ad5-YFV and LMA vaccines in 2-dose (prime-boost) regimens in which Ad5-YFV and LMA were administered 21 days apart in various combinations (Fig 5A). Serum was collected 42 days after the 2^nd^ vaccination as well as 28 days post-infection. F1-V specific IgA was determined by ELISA. Titers were determined in triplicate. One-way ANOVA with Tukey’s post hoc test was used to determine significant differences between groups. Asterisks indicate significance compared to naïve serum. **** p<0.0001.

